# An *in vivo* “turning model” reveals new RanBP9 interactions in lung macrophages

**DOI:** 10.1101/2024.05.22.595416

**Authors:** Yasuko Kajimura, Anna Tessari, Arturo Orlacchio, Alexandra Thoms, Maria Concetta Cufaro, Federica Di Marco, Foued Amari, Min Chen, Shimaa H.A. Soliman, Lara Rizzotto, Liwen Zhang, Joseph Amann, David P. Carbone, Amer Ahmed, Giuseppe Fiermonte, Mike Freitas, Alessia Lodi, Piero Del Boccio, Dario Palmieri, Vincenzo Coppola

## Abstract

The biological functions of the scaffold protein Ran Binding Protein 9 (RanBP9) remain elusive in macrophages or any other cell type where this protein is expressed together with its CTLH (C-terminal to LisH) complex partners. We have engineered a new mouse model, named RanBP9-TurnX, where RanBP9 fused to three copies of the HA tag (RanBP9-3xHA) can be turned into RanBP9-V5 tagged upon Cre-mediated recombination. We created this model to enable stringent biochemical studies at cell type specific level throughout the entire organism. Here, we have used this tool crossed with LysM-Cre transgenic mice to identify RanBP9 interactions in lung macrophages. We show that RanBP9-V5 and RanBP9-3xHA can be both co-immunoprecipitated with the known members of the CTLH complex from the same whole lung lysates. However, more than ninety percent of the proteins pulled down by RanBP9-V5 differ from those pulled-down by RanBP9-HA. The lung RanBP9-V5 associated proteome includes previously unknown interactions with macrophage-specific proteins as well as with players of the innate immune response, DNA damage response, metabolism, and mitochondrial function. This work provides the first lung specific RanBP9-associated interactome in physiological conditions and reveals that RanBP9 and the CTLH complex could be key regulators of macrophage bioenergetics and immune functions.

## Introduction

RanBP9 (Ran Binding Protein 9) is a scaffold protein and a key member of the ubiquitously expressed CTLH (C-terminal to LisH) complex, a multi-subunit E3 ligase of poorly characterized functions ^1–3^. In its canonical configuration, the CTLH complex is a heterodecameric macro-aggregate that includes Armc8 (Armadillo repeat containing 8), Gid4 (Glucose-induced degradation deficient 4), Gid8 (Glucose-induced degradation deficient 8), Maea (Macrophage erythroblast attacher, E3 ubiquitin ligase), Mkln1 (Muskelin 1), RanBP9, RanBP10 (Ran Binding Protein 10), Rmnd5A or Rmnd5B (Required for meiotic nuclear division 5 homolog A or B), Wdr26 (WD repeat domain 26), and Ypel5 (Yippee like 5).

In vitro, RanBP9 is key to the formation of supra-molecular configurations of the CTLH complex ^4^. Current working models indicate that RanBP9 acts as a scaffold and aids the ubiquitination of selected substrates by the CTLH complex ^5, 6^. RanBP9 and the CTLH complex have been implicated in a variety of physiological and pathological states including embryonic development and cancer ^1, 5^. While the absence of RanBP9 causes perinatal lethality, occasional RanBP9-deficient mouse survivors on a mixed genetic background display reduced body size and severe sterility both in males and females ^7^. Expression of RanBP9 increases in several types of malignancies ^1, 8–11^. Due to the weakening of DNA repair in RanBP9-deficient cells, RanBP9 has been put forward as a potential target of therapy in non-small cell lung cancer ^9, 10, 12^. However, RanBP9 physiological functions in lungs are not known. In this regard, most of the knowledge about RanBP9 interactions and functions has been gained by using cancer cells where RanBP9 is generally expressed at levels higher than normal. Those paradigms where RanBP9 is artificially knocked-down, knocked-out, or over-expressed likely do not reflect normal physiology ^1, 5^. In addition, cancer cells can adapt to the inactivation or to expression level variations of the CTLH complex ^13^, in line with the idea that this multi-subunit E3 ligase is a protean macro-aggregate that can change its configuration in response to various types of stress ^14^. RanBP9 and related configurations of the CTLH complex are found in most cell types and they have been shown to be involved in the disparate organismal processes such as maternal-to-zygote transition, development, erythroid maturation, and fertility ^7, 15–18^. At cellular level, evidence indicates that RanBP9 might have major roles in basic processes such as various steps of RNA translation and processing, response to DNA damage, cell adhesion, migration, and metabolism among others ^1, 2, 5, 19, 20^. All considered, it is becoming increasingly clear that the CTLH complex functions are likely cell type- and cell context-dependent. Therefore, it is necessary to study this multi-subunit complex in physiological conditions and in the whole organism context to fully understand its biological functions.

Studying RanBP9 *in vivo* can be challenging due to the constant dynamic interplay between different cell types and the fluctuations of expression that can confound experimental results. An additional challenge is represented by the presence of RanBP10, a highly similar paralog, which makes difficult the identification of RanBP9 *in vivo*. To date, there is no antibody that can specifically identify RanBP10 via immunohistochemistry and immunoprecipitation, and some of the commercially available antibodies against RanBP9 potentially recognize also RanBP10 ^16^ (www.humanproteinatlas.org). To overcome some of these limitations, we had previously engineered a mouse model, called RanBP9-TT, where endogenous RanBP9 bears a double tag (TT) made of both HA and V5 together to unequivocally identify RanBP9 in tissues and organs ^21^. These two small tags were deliberately chosen to avoid potential steric hindrance that could block or alter the protein-protein interactions entertained by RanBP9 with other proteins and especially its participation in the CTLH complex. Most importantly, the RanBP9-TT mouse is viable, fertile, and does not show any overt phenotype ^21^. Furthermore, RanBP9 tagged with HA-V5 co-immunoprecipitates with other members of the CTLH complex indicating that the expected interactions are retained ^21^. All considered, the RanBP9- TT mouse proved to be a valuable tool to perform biochemistry experiments *in vivo* and established that the small tags at the C-terminus did not interfere with RanBP9 biological functions. However, the RanBP9-TT model did not overcome the limitation of identifying RanBP9 in a cell-type specific manner *in vivo*. Therefore, we decided to engineer a new model where two different tags can be switched by Cre-recombination. To this aim, we took advantage of CRISPR/Cas9 targeting strategies to engineer a “turning” system where tags can be changed in a Cre/loxP-dependent manner with in mind the idea that the tool could be coupled with any existing Cre mouse line for tissue/cell type-selective *in vivo* investigations. Due to the feature of changing RanBP9-3xHA into RanBP9-V5, we called this new model “RanBP9-TurnX”, where the X refers to the three copies of HA.

Using this new tool, we can provide the first report of a comprehensive cell type specific RanBP9 interactome in normal cells *in vivo*. In particular, we provide the first RanBP9-associated interactome enriched from lung macrophages motivated by the earlier observation that normal alveolar macrophages and tumor associated macrophages in non-small cell lung cancer express high levels of this protein ^10, 21^. Ultimately, this work establishes the RanBP9-TurnX model as a powerful tool to study the dynamic interactions of RanBP9 and the RanBP9-built CTLH complex in different cell types in the context of the whole organism, and reveals their potential role in regulating macrophage physiology.

## Material and Methods

### Mice

All mouse experiments were conducted in accordance with The Ohio State University Institutional Animal Care and Use Committee (IACUC) guidelines. This study was approved by IACUC (protocol number 2008 A0009-R5 titled “Generation, analysis and training in the use of gene knock-out and transgenic rodents”; Principal Investigator: V. Coppola). C57Bl/6Tac mice were purchased from Taconic Biosciences (Germantown, NY, USA). Lysozyme-Cre mice (LysM-Cre; JAX strain #: 028054) and EIIa-Cre mice (strain #: 003724) were purchased from The Jackson Laboratory (Bar Harbor, ME, USA). Mice were housed in a temperature-controlled room in a 12-hour light/dark cycle with a standard diet and water *ad libitum*.

### Generation and Genotyping of RanBP9-TurnX mouse

RanBP9-TurnX mouse was generated by The Genetically Engineered Mouse Modeling Core of The Ohio State University Comprehensive Cancer Center using CRISPR/Cas9 technology. Briefly, the CRISPR/Cas9 knock-in targeting strategy was designed using the online Benchling software (https://www.benchling.com). The synthetic single strand oligo donor DNA (ssODN) containing the Cre-dependent switchable 3XHA-V5 sequence was purchased from Genewiz LLC, South Plainfield, NJ 07080, USA). The ssODN was 292 base pair long: 5’-GGTCAGGAGTTGGGTCCTGTGCATTTGCCACAGTGGAAGACTACCTACATATAACTTCGTATAGCCTACATTATACGAAGTTATCTTACCCATACGATGTTCCAGATTACGCTTACCCATACGATGTTCCAGATTACGCTTACCCATACGATGTTCCAGATTACGCTTAGCATAACTTCGTATAGCCTACATTATACGAAGTTATCTGGTAAGCCTAT CCCTAACCCTCTCCTCGGTCTCGATTCTACGTAGCTATGCACTTCAAGAGCTCACACTCACATTGTGGCAAACA-3’. Synthetic tracrRNA and crRNA was purchased from Sigma-Aldrich (Saint Louis, MO, USA). GeneArt Platinum Cas9 nuclease protein (cat. nr. B25642) was purchased from Invitrogen (ThermoFisher Scientific; Waltham, MA, USA). The pre-assembled mixture of sgRNA, Cas9 protein, and ssODN was microinjected into C57Bl/6Tac zygotes. C57Bl/6Tac animals were used to propagate colonies from putative founders. Genotyping was determined by PCR analysis of genomic DNA extracted from mouse tail. PCR was performed with Phire Green Hot Start II PCR Master Mix (Thermo Fisher Scientific). Primers used to genotype RanBP9-TurnX mice were: Forward (F) 5’-AAA CCC ACA ATC TGC CAA AG-3’ and Reverse (R) 5’-AAA GCG ACA AAA ACCTGT CC-3’. The WT non-mutated RanBP9 allele produces a fragment of 256 base pairs (bp) while the RanBP9-TurnX allele produces a fragment of 455 bp that is digested into two fragments of 317 and 138 bp after digestion with TaqaI. Following Cre recombination the F and R primers amplify a product of 334 bp, which can be digested into two fragments of 196 and 138 bp by TaqaI. Primers recommended by the Jackson Laboratories were used to identify SpC-Cre^ERT2^ or EIIa-Cre transgene. Sanger sequencing was performed using gel-purified PCR product by QIAquick Gel Extraction Kit (QIAGEN).

### Protein extractions and Western blotting

Mice were sacrificed to excise organs (lung, liver, kidney, spleen, and thymus). Organs were smashed through a 100 mm strainer, followed by lysis of erythrocytes in eBioscience 1X RBC Lysis Buffer (Thermo Fisher Scientific). After washing, proteins were extracted using NP-40 buffer with Halt Protease and Phosphatase Inhibitor Single-Use Cocktail (Thermo Fisher Scientific) and Pepstatin (Roche). Protein concentrations were assessed by Protein Assay Dye Reagent Concentrate (BIO-RAD) and NanoDrop One (Thermo Fisher Scientific). 30 to 50 mg of total protein was run on Mini-PROTEAN TGX Gels (BIO-RAD) and transferred to Nitrocellulose Membranes (BIO-RAD). After blocking with 5% Blotting-Grade Blocker (BIO-RAD) in Tris buffer saline with 0.1% Tween 20 (TBS-T) for 1 hour at room temperature, the membranes were incubated with primary antibodies overnight at 4°C. The proteins of interest were visualized using horseradish peroxidase (HRP)-conjugated secondary antibodies and enhanced by SuperSignal West Pico PLUS Chemiluminescent Substrate (Thermo Fisher Scientific) or SuperSignal West Femto Maximum Sensitivity Substrate (Thermo Fisher Scientific). The antibodies used were anti-RanBP9 (Abcam), anti-HA-tag (Cell Signaling Technology), anti-V5-tag (Invitrogen), HRP-conjugated anti-Rabbit (Abcam), and HRP-conjugated anti-Mouse (Kindle Biosciences) with a dilution in the blocking buffer above. Equivalent loading among samples was confirmed with HRP-conjugated anti-Vinculin (Cell Signaling Technology) or HRP-conjugated anti-GAPDH (Cell Signaling Technology). Western blotting (WB) results were visualized and analyzed by KwikQuant Imager (Kindle Biosciences).

### Immunoprecipitation (IP)

For IP purposes, organs were dissected, and single cells suspension were obtained by using the gentleMACS organ-specific protocol and reagents according to manufacturer’s instructions. Briefly, lungs and livers were treated enzymatically and mechanically by using the Lung Dissociation Kit, mouse (Miltenyi Biotec) and the Liver Dissociation Kit, mouse (Miltenyi Biotec), respectively. Bone marrow-derived macrophages (BMDMs) were differentiated using 20 ng/ml of recombinant murine macrophage colony-stimulating factor (PeproTech) from bone marrow cells which were harvested from femurs and tibias. Erythrocytes were lysed with Red Blood Cell Lysis Solution (Miltenyi Biotec). Cell suspensions were lysed in NP-40 buffer with Halt Protease and Phosphatase Inhibitor Single-Use Cocktail (Thermo Fisher Scientific) and Pepstatin (Roche) to obtain protein lysates. For each IP, 1 mg of total protein was pre-cleared by Pierce Protein A/G Magnetic Beads (Cell Signaling Technology) for 1 hour at room temperature. The pre-cleared lysate was then mixed with anti-HA magnetic beads (Cell Signaling Technology) or anti-V5 magnetic beads (Cell Signaling Technology), and incubated rotating overnight at 4°C. After being washed 5 times in NP-40 buffer, the bound proteins were eluted in Pierce Lane Marker Reducing Sample Buffer (Thermo Fisher Scientific) and boiled for 5 min. Input fractions and corresponding immunoprecipitates were analyzed by WB. The following primary antibodies were used: anti-RanBP9 (Abcam), anti-V5-tag (Invitrogen), anti-GID8 (Proteintech), MAEA (R&D SYSTEMS), Clean-Blot IP Detection Reagent (Thermo Fisher Scientific) was used as a secondary antibody when antibody interferences occurred.

### Immunoprecipitation tandem mass spectrometry (IP-MS/MS)

After harvesting lungs, single cell suspensions were prepared using mouse Lung Dissociation Kit (Miltenyi Biotec) and gentleMACS Octo Dissociator with Heaters (Miltenyi Biotec). Erythrocytes were lysed with Red Blood Cell Lysis Solution (10×) (Miltenyi Biotec). The cell lysis buffer for protein extraction and the reagent to measure protein concentrations are the same as above. Subsequently, 1 mg of pre-cleared protein lysates were bounded with anti-HA magnetic beads (Cell Signaling Technology) or anti-V5 magnetic beads (Cell Signaling Technology) with rotation overnight at 4°C. After washing with the cell lysis buffer stringently, beads were equilibrated in 50 mM of ammonium bicarbonate with 2 washes followed by on-bead digestion with trypsin (800 ng; Promega cat. nr. V5280) overnight at 37 °C, 800 rpm. The digestion continued the next day for 4 hours after samples were supplemented with an additional bolus of trypsin (800 ng). Beads were placed in a magnetic stand to collect supernatants and dried down in a Speedvac concentrator. Prior to mass spectrometry analysis, peptides were resuspended in loading buffer (2% ACN, 0.003% TFA) and quantified via Nanodrop. Chromatography separation and mass spectrometry methods were performed as described in Scheltema *et al*. for data dependent acquisition on a Q-Exactive HF (Thermo Fisher Scientific) ^22^. Tryptic peptides (1000 ng) were isocratically loaded onto a PepMap C18 trap column (300 µm x 5 mm, 100 Å, 5 µm) at 5 µL min−1 and analytical reversed phase separations were performed on a DionexUltiMate 3000 RSLCnano HPLC system coupled to an EASYSprayPepMap C18 column (15 cm × 50 µm ID, 100 Å, 2 µm) over a 90 min gradient at 300 nL min−1. RAW mass spectrometry files were converted to mzml and searched against a complete, reviewed mouse Uniprot database containing common contaminants (downloaded 04/10/2019) via OpenMS (version 2.3.0) with X! TANDEM (release 2015.12.15.2) and MS-GF + (release v2018.01.20).

### Bioinformatics and Proteomics data processing

Proteomics MS/MS raw data were processed using Proteome Discoverer^TM^ v. 2.4 (Thermo Fisher Scientific, US) and Sequest HT search engine against the UniProt database (released 2021_06, taxonomy Mus musculus (Mouse), 55,336 sequences). Processing parameters were set as follows: trypsin digestion was specified as digestion mode with a maximum of two missed cleavages, precursor mass tolerance 10 ppm, and 0.6 Da for the HCD fragment ions. Carbamidomethylation of cysteines (C) was defined as fixed and quantification modification, while oxidation of methionine (M) was set as variable modifications. False discovery rate (FDR) was set to 1% for both protein and for peptide levels. Minora Feature Detection node was used to detect peptide identifications in individual LC-MS/MS runs for subsequent chromatographic alignment to have the same retention time across all sample files (maximum RT shift 10 min, mass tolerance 10 ppm). Protein ratio was calculated using the pairwise ratio approach as the geometric median of the peptide group ratios for each quantified protein. Using this method, we evaluated not only the different protein expression, but also the presence and absence of proteins between the different immunoprecipitates. t-test with a *p*-value threshold of 0.05 was used to define the statistical significance of proteins differentially expressed between the performed comparisons.

## Results

### Generation of the RanBP9-TurnX allele

We used a CRISPR/Cas9 genome editing strategy to knock-in a loxP-3xHA-loxP-V5 cassette before the stop codon at the C-terminus of RanBP9. For targeting purposes, we employed the same guide RNA previously used to generate the RanBP9-TT model (***Figure 1A***, ***Supplementary Figure 1A-B***) ^21^. The single strand oligo DNA (ssODN) to insert the HA-V5 turning element at the 3’ end of RanBP9 was 292 bp and contained three copies of the HA tag (3xHA) in order to maintain a distance between the two loxP sites sufficient for the Cre enzyme to excise the intervening sequence. A minimal linker sequence was inserted downstream the 3’ loxP site to keep the successive V5 tag in frame. Both the 3xHA and the V5 sequence included a STOP codon at the 3’ end (***Figure 1A***, ***Supplementary Figure 1A,C***). C57Bl/6Tac zygote microinjections with CRISPR/Cas9 reagents produced three potential founders (F0) bearing the confirmed insertion of the loxP-3xHA-loxP-V5 cassette at least on one of the two RanBP9 alleles (Females #19 and #35, and male #32). All the three founders were backcrossed to WT C57Bl/6Tac mice. Male founder #32 was the first to produce heterozygous progeny (F1) positive for the correct insertion of the “turning cassette” and this line was chosen for further propagation. F1 heterozygous mice were crossed to both each other and to WT C57Bl/6Tac mice for further backcrossing. PCR screening of F2 homozygous animals showed the expected RanBP9-TurnX band (455 bp) and Sanger sequencing confirmed that the loxP-3xHA^STOP^-loxP-V5^STOP^ was successfully inserted at the C-terminus of RanBP9 (***Figure 1B***, ***Supplementary Figure 1D-E***). RanBP9-TurnX heterozygous parents produced the expected number of RanBP9-TurnX homozygous (from now on indicated as BP9HA mice; see ***Table 1***), both male and female pups in the absence of any apparent abnormal phenotype. Like the RanBP9-TT mice that we had previously engineered ^21^, BP9HA mice are viable and fertile and do not show any overt phenotype (***Supplementary Figure 1F***). Likewise, organs and body size compared to WT control mice by basic observational and anatomical examination (not shown). These results indicated that the RanBP9-TurnX allele does not cause lethality and it is producing a fully functional RanBP9 tagged protein. See ***Table 1*** for the nomenclature used to indicate mouse strains, proteins produced, and cell types where the indicated proteins are expressed.

**Table 1.** List of mouse models, wild type (WT) and tagged RanBP9 protein expression. In RanBP9-TurnX homozygous animals (**BP9HA** mice), RanBP9 is expressed as a fusion product of WT RanBP9 with 3 copies of an HA tag at the C-terminus. In EIIa-Cre::RanBP9-TurnX homozygous animal (**BP9V5** mice), RanBP9 is “turned” into a fusion product of WT RanBP9 with a V5 tag at the C-terminus in all cell types. In LysM-Cre::RanBP9-TurnX homozygous animals (LysMBP9X), cells not expressing Cre (LysM-Cre^NEG^) will continue to express RanBP9 tagged with HA while LysM-Cre positive cells (macrophage-enriched) will express RanBP9 turned into RanBP9 with a V5 tag at the C-terminus. Control C57Bl/6-Taconic, EIIa-Cre, and LysM-Cre mice express RanBP9 as WT protein. Nomenclature of mouse strains generated and used for this work.

| MOUSE STRAIN NAME | ABBREVIATION | TAG | RanBP9 PROTEIN PRODUCT | RANBP9 CELL TYPE EXPRESSION |
| --- | --- | --- | --- | --- |
| C57Bl/6-Taconic | B6 WT | NONE | RanBP9 WT | UBIQUITOUS |
| RanBP9-Turnx homozygous | BP9HA | 3xHA | RanBP9-3xHA | UBIQUITOUS |
| Ella-Cre | E2aBP9 WT | NONE | RanBP9 WT | UBIQUITOUS |
| Ella-Cre::RanBP9-Turnx homozygous | E2aBP9V5 | V5 | RanBP9-V5 | UBIQUITOUS |
| LysM-Cre | LysMBP9 WT | NONE | RanBP9 WT | UBIQUITOUS |
| LysM-Cre::RanBP9-Turnx homozygous | LysMBP9X | 3xHA<br>V5 | RanBP9-3xHA<br>RanBP9-V5 | LysM-Cre<sup>NEG</sup><br>(LysM-Cre negative)<br>LysM-Cre<sup>POS</sup><br>(LysM-Cre positive)<br>(macrophage-enriched) |

**Figure 1.**
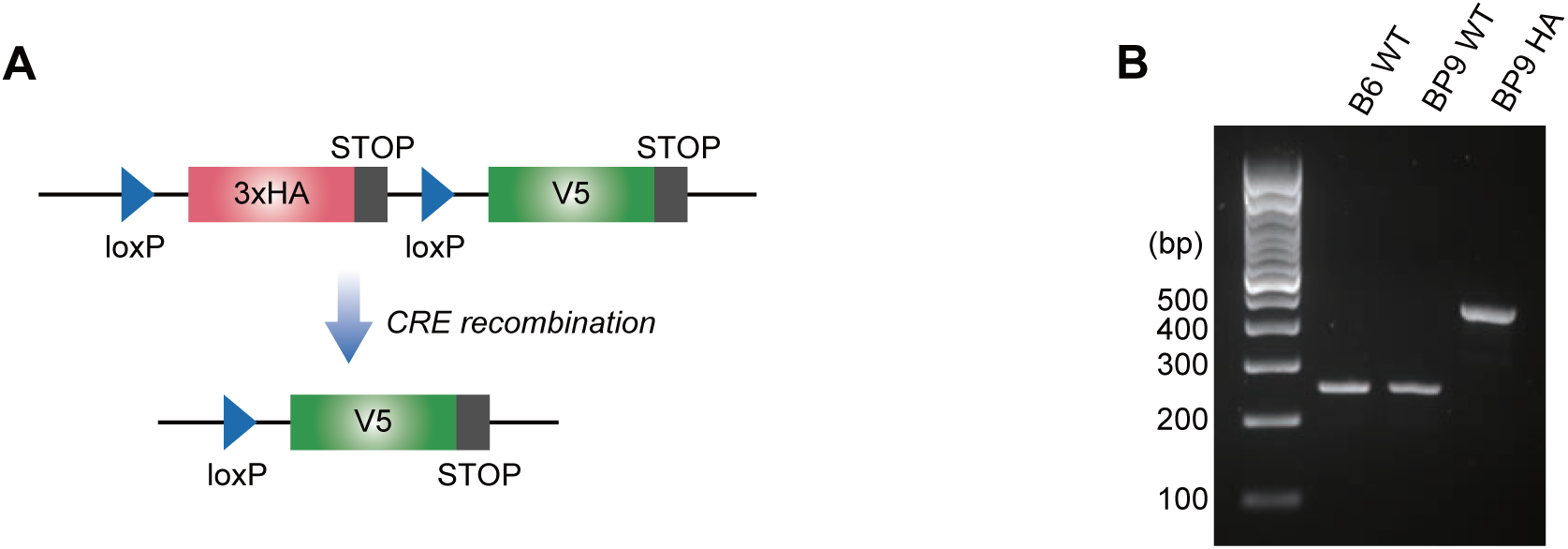
Generation of the RanBP9-TurnX allele. **A**) The RanBP9-TurnX allele includes 3 copies (3x) of the HA tag and a stop codon at the C-terminus of the third HA copy flanked by loxP sites. Cre recombination eliminates the 3xHA cassette and licenses the expression of a V5 tag (followed by a stop codon). **B**) Genotyping of RanBP9-TurnX homozygous mice shows the presence of the expected 455 bp knock-in product.

### Cre-Lox recombination in RanBP9-TurnX mice turns RanBP9-3xHA into RanBP9-V5

For an organism-wide functional validation of the RanBP9-TurnX allele, we crossed two RanBP9-TurnX heterozygous F1 animals with homozygous EIIa-Cre mice commonly used as Cre deleter (indicated as E2A-Cre; JAX strain #:003724) (***Table 1***, ***Figure 2A***). The first generation of progeny from this cross showed that heterozygous “turned” mice were born as expected and indistinguishable from E2aBP9 WT littermates. Then, we crossed heterozygous “turned” animals to each other to obtain homozygous “turned” animals (E2aBP9V5 mice) and littermate controls. The PCR screening amplified from of E2aBP9V5 homozygous “turned” mice one product of the expected 334 bp size (***Figure 2B***). Sanger sequencing confirmed the presence of the loxP-V5^STOP^ sequence and the absence of the loxP-3xHA^STOP^ sequence (***Supplementary Figure 2A***). E2aBP9V5 mice are viable and there are not obvious differences in organs, body size, and the number of females and males born compared to E2aBP9 WT mice. Moreover, E2aBP9V5 “turned” homozygous mice are fertile and do not display any significant body size difference compared to E2aBP9 WT controls (***Supplementary Figure 2B, 2C***).

**Figure 2.**
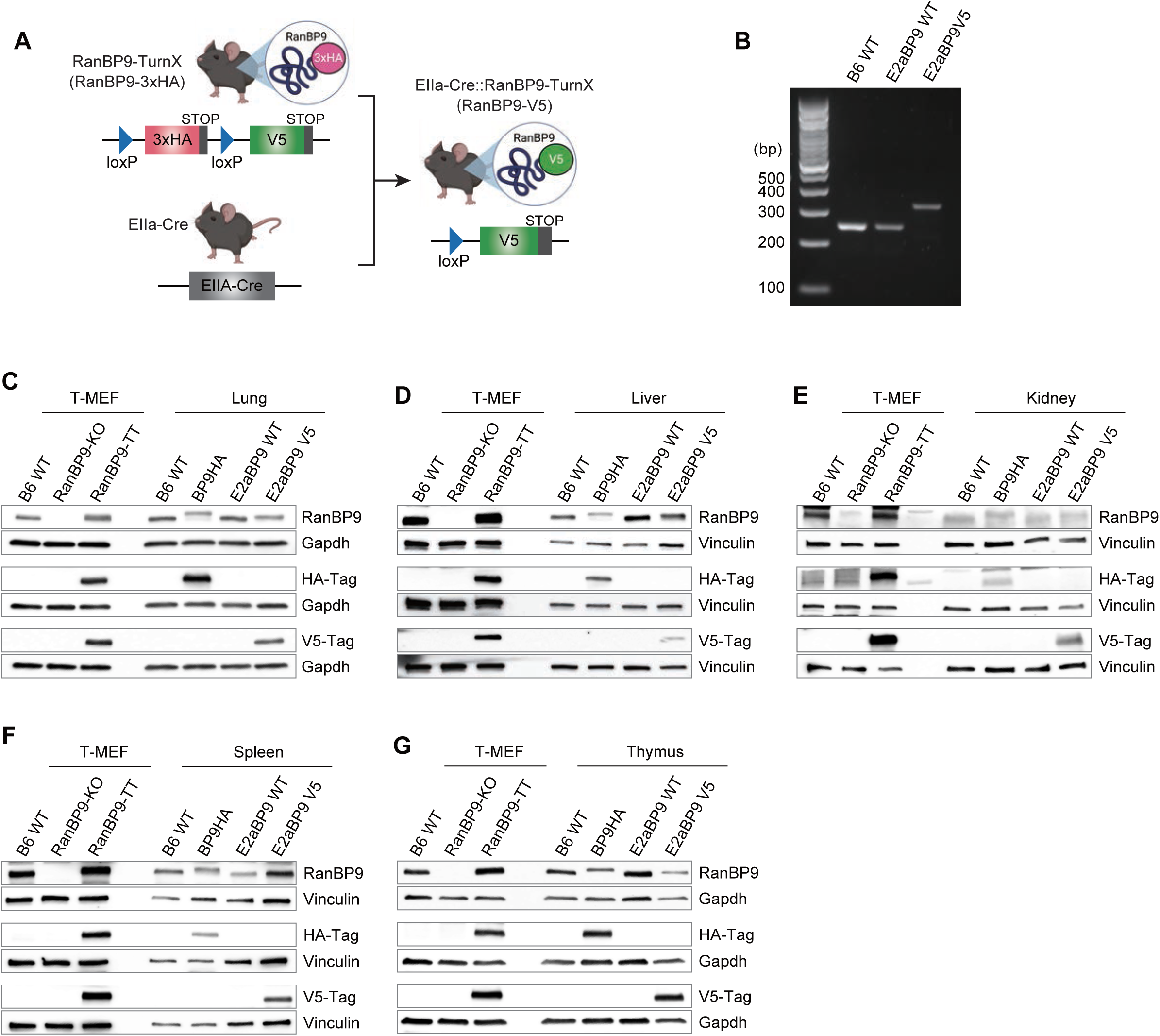
Cre-lox recombination in RanBP9-TurnX mice turns RanBP9-3xHA into RanBP9-V5. **A**) Schematic representation of the “turning from RanBP9-3xHA to RanBP9-V5 expression upon Cre-lox recombination. loxP site sequences work as linkers between RanBP9 and the two different tags. **B**) Genotyping by PCR amplification with the indicated primers produces the expected 334 bp fragment in RanBP9-V5 mice. **C**-**G**) WB analysis of lung (C), liver (D), kidney (E), spleen (F), and thymus (G) performed with the indicated antibodies. WT, RanBP9 knock-out (KO), and RanBP9-TT Mouse Embryonic Fibroblast immortalized with the large-T antigen (T-MEFs) ^21^ were used as positive and negative controls for RanBP9 WT protein and RanBP9-HA or RanBP9-V5 tagged proteins. RanBP9-3xHA is present only in RanBP9-TurnX homozygous animals while RanBP9-V5 is only detected in RanBP9-V5 turned homozygous animals where the EIIa-Cre is also present. Gapdh and Vinculin are used as loading controls.

Next, we sought to ascertain the RanBP9 protein expression both in BP9HA and BP9V5 homozygous “turned” animals. The Western Blot (WB) analysis of lysates probed with anti-RanBP9 antibody showed that RanBP9 in BP9HA mice was clearly detectable in all the organs we analyzed, which included lung, liver, kidney, spleen, and thymus (***Figure 2C-G***). RanBP9-3xHA fusion protein was detected using the HA-specific antibody in the lysates of BP9HA organs, while the V5-specific antibody did not detect any RanBP9-V5 fusion protein (***Figure 2C-G***). These results confirmed that the 3xHA-tagged RanBP9 fusion protein in BP9HA mice is produced and detected as expected. On the other hand, BP9V5 was clearly detected only in organs from turned animals (***Figure 2C-G***). Most importantly, turned animals did not show any detectable RanBP9-3xHA protein. Collectively, these results showed that RanBP9 tag is switched from 3xHA to V5 upon Cre recombination as expected without any functional consequence on RanBP9 protein expression.

### RanBP9-V5 can be efficiently immunoprecipitated from BP9V5 mouse organs

Next, we sought to determine whether it was possible to immunoprecipitate the V5-tagged RanBP9 protein from RanBP9-turned organs. It was expected that RanBP9 co-immunoprecipitated with known members of the CTLH complex. We performed immunoprecipitations from lung, liver, and BMDM, and tested for the presence of Gid8 and Maea, two core components of the CTLH complex directly interacting with RanBP9 (***Figure 3***) ^5^. WB results clearly showed the successful immunoprecipitation of these two RanBP9 binding partners.

**Figure 3.**
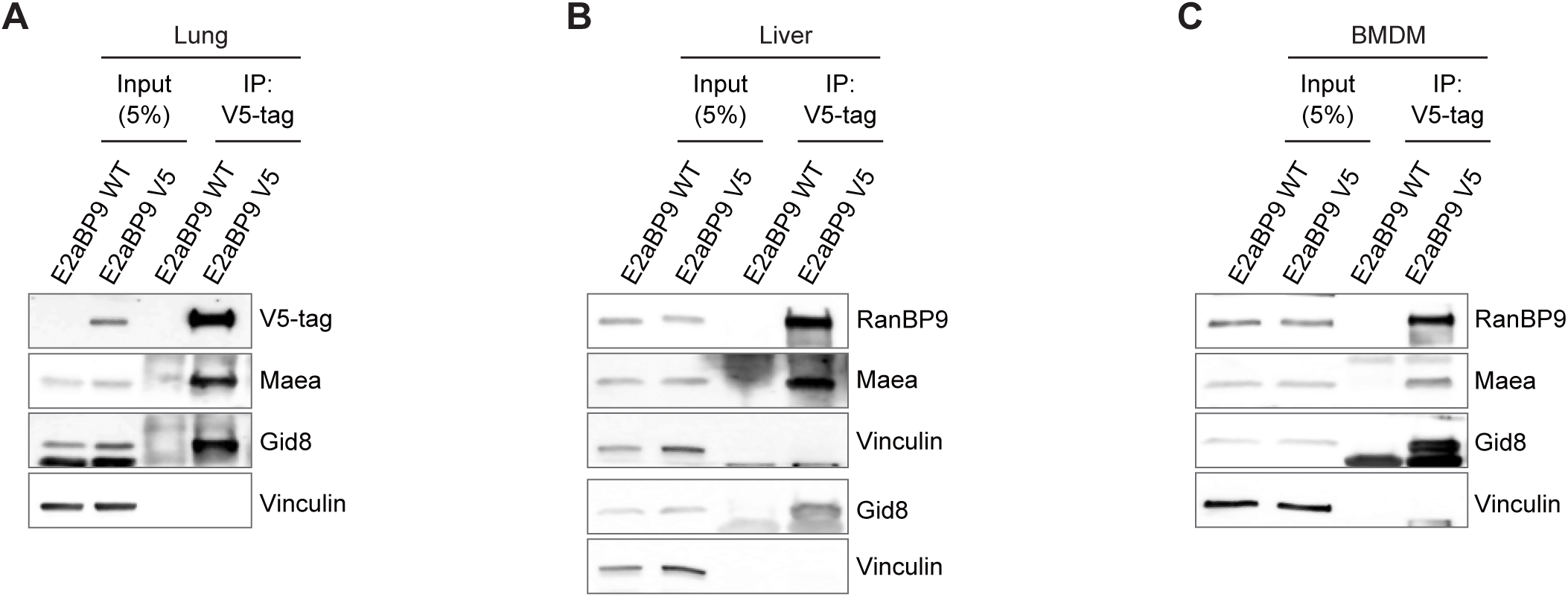
RanBP9-V5 can be efficiently immunoprecipitated from RanBP9 “turned” mouse organs. Immunoprecipitation using anti-V5 conjugated beads shows the successful pull-down of RanBP9-V5 from A) lung, B) liver, and C) bone marrow derived macrophages (BMDM). In all three organs, RanBP9 co-immunoprecipitates with the two CTLH members and binding partners Gid8 and Maea. Vinculin is used as loading control.

These results unequivocally showed that RanBP9-V5 is produced as expected upon recombination in all cells expressing this protein. Furthermore, RanBP9-V5 fusion protein takes part in the formation of the CTLH complex.

### LysM-Cre-mediated recombination licenses the expression of RanBP9-V5 in lung

RanBP9 is expressed by both normal lung epithelial cells and alveolar macrophages ^10, 21^. However, very little is known about RanBP9 and the CTLH complex in those specific cell types. In this regard, one of the core CTLH proteins, Maea, has been shown to have a role in macrophages involved in reticulocyte enucleation ^23^. To generate a model where RanBP9 in macrophages bore a different tag in comparison to all the other cell types, we crossed RanBP9-TurnX mice with the widely used strain B6.129P2-Lyz2^tm1(cre)Ifo^/J (JAX strain #: 028054), also known as LysM-Cre mice ^24^. In this model, LysM-positive (LysM^POS^) cells, including resident alveolar macrophages, express RanBP9-V5 while other non-recombinant cells maintain the expression of RanBP9 tagged with 3xHA. Our goal was to identify the interactions entertained by RanBP9 in lung macrophages and the LysM-Cre mouse targets nearly all the resident lung macrophages including the alveolar population, which we had found highly positive for RanBP9 ^10, 21, 25^. The WB analysis of the whole lung lysates showed the presence of both RanBP9-3xHA and RanBP9-V5 (***Figure 4***). These results demonstrate that RanBP9-V5 expression licensed by LysM-Cre is clearly detectable in LysMBP9X mice.

**Figure 4.**
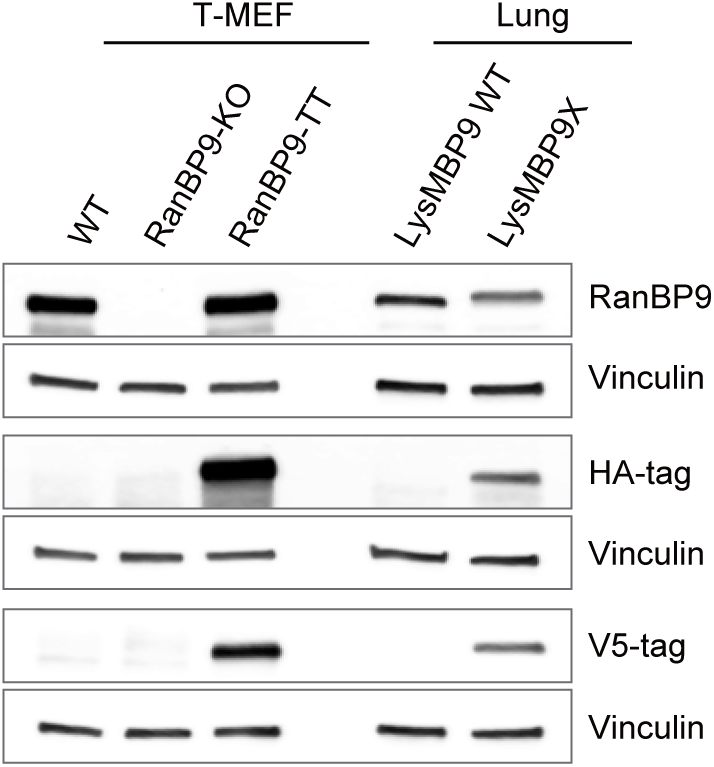
LysM-Cre licenses the expression of RanBP9-V5 in lung. WB analysis shows that in LysMBP9X animals both RanBP9-3xHA and RanBP9-V5 are clearly detectable while they are absent in LysMBP9 WT controls. WT, RanBP9 KO, RanBP9-TT T-MEFs were used as positive and negative controls. Vinculin is used as loading control.

### The RanBP9-TurnX model allows the efficient immunoprecipitation of the CTLH complex in normal lungs both by HA and V5 tags

Next, we took one LysMBP9X and one LysMBP9 WT control, obtained lysates from single cell suspensions, and performed immunoprecipitation using HA-, V5-, and Ig control-conjugated beads. Immunoprecipitated fractions were stored at −80°C until mass spectrometry (MS) data acquisition of all samples was performed. We repeated this process four times for a total of four independent lysates from LysMBP9X lungs and four independent LysMBP9 WT controls. Finally, all the 24 immunoprecipitated fractions were analyzed by MS. The negative controls of the immunoprecipitated fractions obtained from WT mice or with isotype control-conjugated beads allowed the exclusion of proteins non-specifically bound to beads and the anti-tag antibodies all representing probable false interactors (***Supplementary Tables 1-4***). A total of 278 proteins remained after eliminating all entries present in the list of HA-immunoprecipitated proteins that did not show a statistical significance of p ≤ 0.05 in comparison with 3 negative control groups (HA-immunoprecipitated from WT, Ig-immunoprecipitated from LysMBP9X mice, and Ig-immunoprecipitated from LysMBP9 WT control mice), (***Supplementary Tables 1****-**2**)*. Eleven (11) of those entries were expected CTLH complex members, clearly indicating that the immunoprecipitation by 3xHA worked efficiently (***Table 2***). We repeated the same purging process for the V5-immunoprecipitated proteomic list in comparison with the three negative control groups (V5-immunoprecipitated from WT, Ig-immunoprecipitated from LysMBP9X mice, and Ig-immunoprecipitated from LysMBP9 WT controls). Ultimately, 202 proteins were significantly enriched (p ≤ 0.05) in the V5-immunoprecipitated fraction (***Supplementary Tables 3-4***). Ten (10) of those proteins were expected CTLH complex members (***Table 3***). Notably, Rmnd5b did not appear in the list of the V5-interactome. Also, in both HA- and V5 RanBP9-interactomes, Wdr26 was identified as two different isoforms (UniProt ID: E0CYH4 of 625 amino acids; and UniProt ID: A0A494BB75 of 725 amino acids). These results clearly demonstrate that both the 3xHA- and the V5-immunoprecipitation was successful in enriching for RanBP9 interactors from whole lung lysates. Also, both RanBP9-3xHA and RanBP9-V5 partake as expected in the CTLH complex in lung protein lysates from LysMBP9X animals.

**Table 2.**
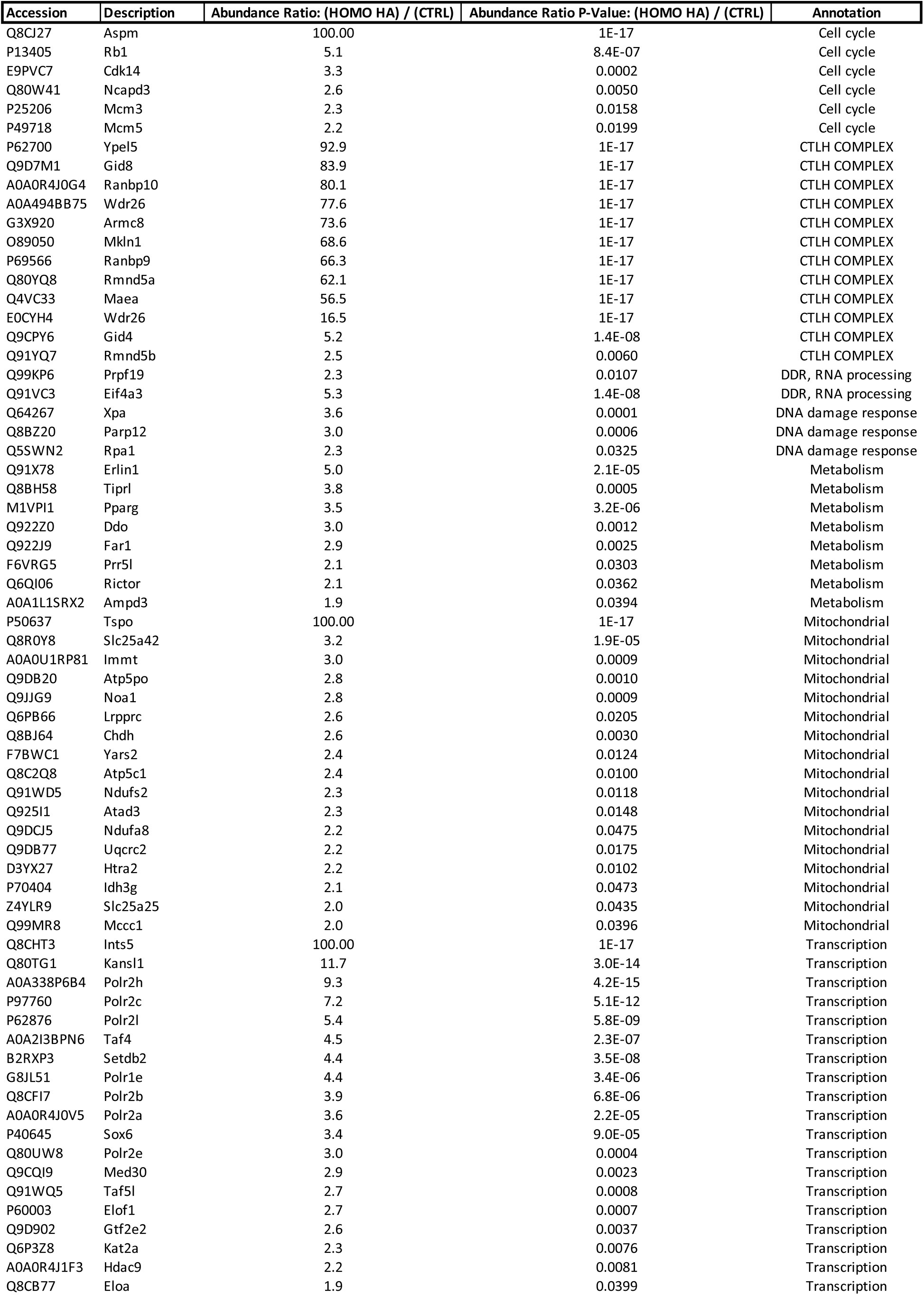
List of selected significant interactors immunoprecipiatated by RanBP9-3xHA.

**Table 3.**
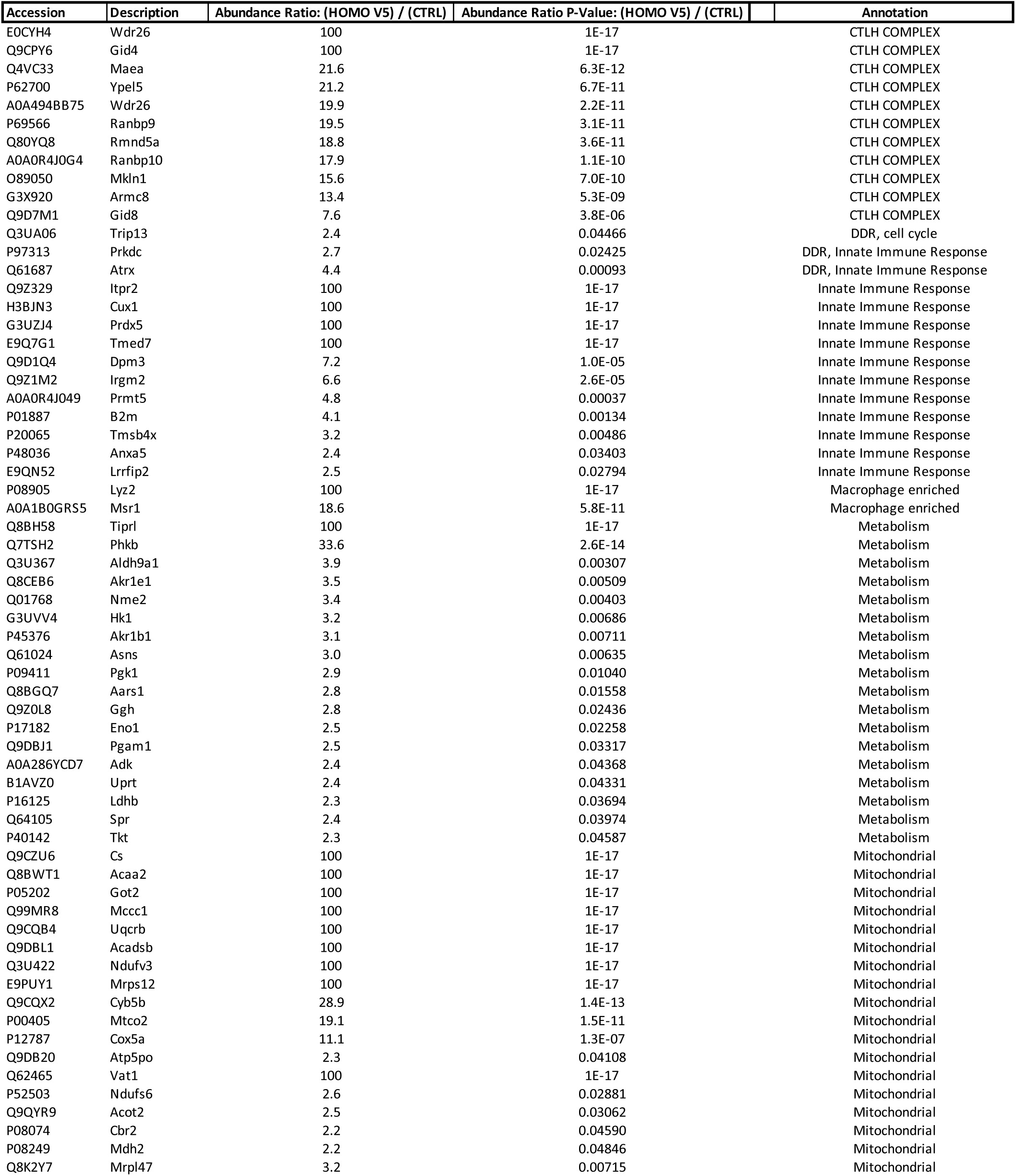
List of selected significant interactors immunoprecipiatated by RanBP9-V5.

### RanBP9-3xHA and the RanBP9-V5 immunoprecipitated proteomes show remarkable differences and include previously unknown interactors

Next, we compared the list of HA-immunoprecipitated proteins with the list of V5-immunoprecipitated ones. Seventeen (17) proteins were common between the HA-immunoprecipitated and the V5-immunoprecipitated proteome, excluding the CTLH complex members immunoprecipitated both by RanBP9-3xHA (11) and RanBP9-V5 (10) (***Supplementary Table 5***). Of the seventeen (17) proteins in common between the 3xHA- and V5-interactome, two were proteins present in the mitochondria and the rest were proteins involved in the most disparate biological processes, or not well characterized (***Supplementary Table 5***).

Strikingly, more than 90% of the proteins immunoprecipitated with the two tags from the same lung lysates were different, being present only in the 3xHA- or the V5-associated list (***Table 2***, ***Table 3***, ***Supplementary Table 2***, ***Supplementary Table 4***). These results strongly indicate that the HA-immunoprecipitated and the V5-immunoprecipitated proteomes are highly enriched for the LysM-Cre^NEG^ and LysM-Cre^POS^ cells, respectively.

Next, we sought to determine how many of the proteins immunoprecipitated by RanBP9-3xHA and RanBP9-V5 had previously been reported (***Supplementary Table 6***). Interestingly, the proteomic lists immunoprecipitated with both the HA and the V5 tag included a majority of RanBP9-interactors not previously reported. Indeed, only 28.4% of the entries from the HA list (***Supplementary Table 7***) and 35.1% of the proteins from the V5 list were previously reported interactors (***Supplementary Table 8***). These results strongly indicate that RanBP9 and the CTLH complex have cell-type specific interactions and functions.

### The lung RanBP9-immunoprecipitated interactome suggests the involvement of the CTLH complex in the regulation of fundamental biological processes in lung cells

To begin to determine what biological processes and cellular pathways might be regulated by RanBP9 and the CTLH complex in lung cells, we performed a Metascape analysis (www.metascape.org) ^26^. First, we performed an analysis that includes all the proteins immunoprecipitated with both HA and V5 tags (***Figure 5***). Using 450 unique entries (446 recognized by the Metascape database), the most significant enriched term was “processing of capped intron containing pre-mRNA” (R-MMU-72203). In line with previously published results, terms such as “mRNA splicing” (R-MMU-72165), “regulation of RNA splicing” (GO:0043484), “alternative mRNA splicing via spliceosome” (GO:000380) were also significantly enriched ^19, 20^. Other significantly enriched terms present in the list indicated the involvement of RanBP9 and the CTLH complex in processes that are in line with previously reported data such as “transcription” ^27, 28^, “HSP90 signaling in connection with steroid hormones” ^29^, “Rho GTPases” ^30^, “mitotic cell cycle” ^31–33^, and the “cytoskeleton” ^5, 34, 35^. Interestingly though, there were also terms that suggest the involvement of RanBP9 and the CTLH complex in “Huntington disease”, “translation”, “glyceraldehyde-3-phosphate biosynthetic process”, “adaptive innate immunity”, and “positive regulation of phagocytosis”.

**Figure 5.** Gene ontology cluster enrichment of comprehensive RanBP9 interactors in mouse lung lysates. 446 unique proteins (of the total 450 RanBP9 putative interactors found in this study) were recognized by the Metascape algorithm and used for a gene set enrichment analysis (www.metascape.org) ^26^.

These results indicate that in normal lungs RanBP9 and the CTLH complex connect directly or indirectly with proteins involved in a variety of fundamental biological processes.

### Analysis of the RanBP9-HA immunoprecipitated proteome

Next, we performed a Metascape analysis of the RanBP9-3xHA immunoprecipitated interactome (***Figure 6***). The list of 275 RanBP9-3xHA interactors recognized by Metascape (out of the 278 total) included proteins involved in RNA-processing, protein-DNA complex organization, negative regulation of small molecule metabolic process, and regulation of receptor mediated endocytosis among others. In regard to RNA splicing, the RanBP9-3xHA-associated interactome included Eif4a3 (Eukaryotic Translation Initiation Factor 4a3), Cwc22 (CWC22 Spliceosome Associated Protein Homolog), Rbm8 (RNA binding motif protein 8), and Magohb (MAGO homolog b) that form the Exon Junction Complex. It also included two members of the THO complex Tho2 and Tho6, and two members of the PRP19C/Prp19 (Pre-mRNA processing factor 19) complex/NTC/Nineteen complex such as Prpf19 and Plrg1 (Pleiotropic regulator 1). Finally, several small nuclear ribonucleoproteins (Snrps) of the Snrp200, Snrp40, including the SMN-Sm-complex and Snrpe, another Pre-m-RNA processing factor were also present (***Table 2***, ***Supplementary Table 2***).

**Figure 6.** Gene ontology cluster enrichment of RanBP9-3xHA interactors in mouse lung lysates. 275 unique proteins (of the total 278) RanBP9-3xHA putative interactors were recognized by the Metascape algorithm and used for a gene set enrichment analysis (www.metascape.org) ^26^

In regard to transcription, in addition to Med30 (Mediator 30), Gtf2e2 (General transcription factor 2e2), Ints5 (Integrator complex subunit 5), Taf4 (TATA-box binding associated factor 4), and Taf5l (TATA-box binding associated factor 5 like), six (6) different subunits of the DNA-directed RNA polymerase II were present in the HA-immunoprecipitated proteome. Additional genes that regulate transcription were also found in this list: Kansl1 (Kat8 regulatory NSL complex subunit 1), Setdb2 (SET domain bifurcated histone lysine methyltransferase 2), Sox6 (SRY-related HMG-box 2), Elof1 (Elongator factor 1), Eloa (Elongin a), Kat2a (Lysine acetyltransferase 2), and Hdac9 (Histone deacetylase 9) (***Table 2***, ***Supplementary Table 2***).

RanBP9 has been shown to regulate cell cycle ^13, 31^. The HA-interactome contained Aspm (Assembly factor for spindle microtubules 3), Ncapd3 (Non-SMC condensing II complex subunit d3), Rb1 (Retinoblastoma-associated protein 1), Cdk14 (Cyclin-dependent kinase 14), and the two members of the replisome Mcm3 (Minichromosome maintenance complex component 3) and Mcm5 (Minichromosome maintenance complex component 3) (***Table 2***, ***Supplementary Table 2***).

We have previously shown that RanBP9 plays a role in the DNA damage response (DDR) and ATM-p53 signaling ^9, 10, 12^. Interestingly, Tiprl (Tip41-like protein) that promotes H2AX phosphorylation and may play a role in ATM signaling ^36, 37^, was pulled-down both by RanBP9-3xHA and RanBP9-V5 (***Table 2***, ***Table 3***).

RanBP9-3xHA coimmunoprecipitated with the already mentioned Eif4a3 that is a helicase of the exon junction complex and participates in the clearing of excessive R-loops ^38, 39^. Furthermore, the RanBP9-3xHA interactome included well-known players of the DDR such as Xpa (Xeroderma pigmentosum group a-complementing protein) that has a central role in nucleotide excision repair ^40^, and Rpa1 (Replication Protein A1) ^41^. RanBP9-3xHA pulled down also Prpf19 (Pre-mRNA processing factor 19) that is recruited to sites of DNA damage and ubiquitinate the RPA complex (Rpa1 and Rpa2), which in turn allows the recruitment and activation of ATR ^42^. Interestingly, RanBP9-3xHA pulled down Parp12 (Poly[ADP-ribose] polymerase member 12), which had been previously reported as RanBP9 interactor ^43^, and is an interferon stimulated gene that contributes to suppressing viral infections (***Table 2***, ***Supplementary Table 2***) ^44, 45^.

Recently, RanBP9 has been reported to regulate glycolysis ^43^. An enzyme involved in carbohydrate metabolism immunoprecipitated by RanBP9-3xHA was Man2c1 (Mannosidase Alpha Class 2C member 1). However, the list of the RanBP9-3xHA interactors included Ampd3 (Adenosine Monophosphate Deaminase 3) ^46^, Entpd1 (Ectonucleoside Triphosphate Diphosphohydrolase 1, also known as CD39) that are involved in nucleotide metabolism, Ddo (D-Aspartate Oxidase) ^47^, which relates to amino acid metabolism, and several proteins regulating lipid metabolism such as Erlin (ER lipid Raft Associated 1), Far1 (Fatty Co-acyl-CoA Reductase 1), Pparg (Peroxisome Proliferator Activated Receptor Gamma). Notably, Rictor (RPTOR independent companion of MTOR complex 2), a major regulator of the MTOR (Mechanistic Target of Rapamycin) pathway and previously reported RanBP9 interactor ^48^ and Prr5l (Proline Rich 5 Like), which is another member of the TOR complex 2, were both identified in the RanBP9-3xHA interactome (***Table 2***, ***Supplementary Table 2***).

These results strongly indicate that the CTLH complex might be involved in a more pervasive regulation of cell metabolism than previously recognized. Remarkably, a total of seventeen (17) proteins with known mitochondrial localization were found in the RanBP9-3xHA-associated proteome (***Table 2***, ***Supplementary Table 2***) suggesting a potential localization of RanBP9 at the mitochondria in physiological conditions. Among the mitochondrial hits it is worth to mention the High temperature requirement protein a2 (Htra2), which has been previously immunoprecipitated with the CTLH complex in different studies ^13, 21, 48^.

### The RanBP9-V5 interactome is enriched for proteins with known functions in macrophages

Finally, we analyzed the RanBP9-V5 immunoprecipitated interactome. The 198 entries recognized by Metascape (out of 202) of the V5-enriched fraction included proteins involved in RNA splicing, amino acid metabolism, signaling by MET (Hepatocyte Growth Factor Receptor), but also energy derivation by oxidation, antigen processing and presentation, IL17A signaling, and defense response to Gram-negative bacteria (***Figure 7***).

**Figure 7.** Gene ontology cluster enrichment of RanBP9-V5 interactors in mouse lung lysates. 198 unique proteins (of the total 202) RanBP9-V5 putative interactors were recognized by the Metascape algorithm and used for a gene set enrichment analysis (www.metascape.org) ^26^

Due to the selective expression of RanBP9-V5 in the LysM-Cre^POS^ cell populations, it was expected to observe not only a different list of interactors compared to the RanBP9-3xHA immunoprecipitated list, but also an enrichment for proteins expressed in the myeloid/monocytic cell fraction. Notably, the RanBP9-V5 interactome was enriched for proteins with established roles in macrophages, players of innate immune response, of cell metabolism, and with mitochondrial localization and functions (***Table 3***, ***Supplementary Table 4***). In this regard, at least thirteen (13) entries have reported roles in macrophage development and function (***Table 3***, ***Supplementary Table 4***).

In particular, we found Lyz2 (Lysozyme 2 or Lysozyme M) and Msr1 (Macrophage scavenger Receptor 1 also known as CD204) that display an enriched expression in monocytes/macrophages. Lyz2 locus drives of the Cre expression in the LysM-Cre strain ^24^. Msr1 is critical to macrophage functions and it was shown to have a role in polarization ^49^. On one hand, their presence is an indication of the successful enrichment of immunoprecipitated proteins from LysM-Cre^POS^ cells. On the other hand, since they were not reported before, these results again support the idea that RanBP9 and the CTLH complex have cell-type specific roles in lungs.

Other proteins that co-immunoprecipitated with RanBP9-V5 and have established roles in macrophages were Anxa5 (Annexin A5), B2m (Beta-2 microglobulin), Cux1 (Cut Like Homeobox 1), Prdx5 (Peroxiredoxin 5), and Tmed7 (Transmembrane P24 Trafficking Protein 7), and Lrrfip2 (Leucine-Rich Repeat Flightless Interacting Protein 2). Anxa5, which was previously reported as a putative RanBP9 interactor ^13^, has major roles in macrophage function reprogramming metabolism through PKM2, regulating inflammatory signals, binding apoptotic cells, and regulating movement.

Its loss specifically reduces the lipopolysaccharide-induced inflammatory response of rodent alveolar macrophages ^50–54^. Specifically in macrophages, β2-microglobulin alters inflammation osteoarthritis ^55^, triggers the inflammasome activation in tumor-associated macrophages ^56^, and determines the response to immunotherapy in lung adenocarcinoma ^57^. Cux1 is involved in the polarization of tumor-associated macrophages by antagonizing NF-κB signaling ^58^. Prdx5 deficiency in macrophages promotes lung cancer progression and M2-like polarization ^59^. Tmed7 is involved in the delivery of TLR4 to the plasma membrane ^60^ and can inhibit TLR4 signaling from the endosome upon LPS stimulation ^61^. Lrrfip2 is also involved in TLR4 signaling and can down-regulate the NLRP3 inflammasome activation ^62, 63^.

RanBP9-V5 also pulled down two major players of the DDR, Prkdc (Protein Kinase, DNA-activated, Catalytic Subunit) better known as DNA-PK, and Atrx (Alpha thalassemia/mental retardation syndrome X-linked). Prkdc is the third of the Apical kinases of the DDR together with ATM and ATR ^64^. Finally, Atrx is involved in replication stress, homologous recombination, and non-homologous end-joining ^65^. Loss of Atrx confers sensitivity to PARP inhibitors similarly to the loss of RanBP9 ^10, 12, 66^.

However, Prkdc and Atrx are also involved in sensing the aberrant presence of DNA in the cytoplasm and, as such, they are major players of the innate immunity response ^67, 68^.

Another noticeable enrichment in the RanBP9-V5 proteome was for enzymes involved in cell metabolism. It is interesting to note that enzymes directly involved in glycolysis like Eno1 (Enolase 1), Hk1 (Hexokinase 1), Ldhb (Lactate dehydrogenase b), Pgam1 (Phosphoglycerate mutase 1), Phkb (Phosphorylase kinase regulatory subunit b), Pgk1 (Phosphoglycerate kinase 1) and Tkt (Transketolase), were significantly enriched only in the LysM-Cre^POS^ (RanBP9-V5) fraction indicating that the CTLH complex may regulate normal lung macrophages glucose utilization, which is central to macrophage polarization and function ^69, 70^.

However, RanBP9-V5 also co-immunoprecipitated with Aars1 (Alanyl-tRNA Synthetase 1), Asns (Asparagine Synthetase), Qdpr (Quinoid Dihydropteridine Reductase), and Ggh (Gamma-Glutamyl Hydrolase) all related to amino acid metabolism; Adk (Adenosine Kinase), Nme2 (Nme/Nme23 Nucleoside Diphosphate Kinase 2), and Uprt (Uracil Phosphoribosyl transferase Homolog) related to nucleotide metabolism; Akr1b1 (Aldo-Keto Reductase Family 1 Member B), Akr1e1 (Aldo-Keto Reductase Family 1 Member E), Aldh9a1 (Aldehyde Dehydrogenase 9 Family Member A1) all related to aldehyde metabolism (***Table 3***, ***Supplementary Table 4***). These results indicate that RanBP9 interacts with enzymes involved in metabolism that is not limited only to glycolysis.

Finally, eighteen (18) entries present in the RanBP9-V5 interactome are proteins that are known to localize to the mitochondria. Notably though, only Atp5po (ATP synthase peripheral stalk subunit OSCP) and Mccc1 (Methylcrotonyl-CoA Carboxyalse subunit 1) were present also in the RanBP9-3xHA list.

These results clearly show that the immunoprecipitation by the V5 tag was successful in enriching for interactors that are expressed in LysM-Cre^POS^ cells and reveal new connections between RanBP9 innate immunity, metabolism, and mitochondria.

## Discussion

RanBP9 is considered an essential member of the CTLH complex, which is a multi-subunit evolutionarily conserved E3 ligase ^1, 2, 5^. Remarkable progress has been made on the topology and the structure of the complex, which can assume different shapes ^6, 14^. In this regard, RanBP9 enables the formation of pro-proliferative supramolecular configurations ^13^. Although the mechanisms are not known, evidence suggests that this complex can perform ubiquitination resulting in the modulation of the functions of the targets without inducing their degradation ^5, 43^.

However, we have only begun to scratch the surface of the biological functions of this fascinating molecular machinery, and it is becoming increasingly clear that, even if it is ubiquitously expressed, the CTLH complex exerts cell type-specific functions. This is exemplified by its involvement in quite different processes like maternal-to-zygote transition and erythroid maturation ^16–18^. Therefore, innovative tools are needed to elucidate how the CTLH complex works in a cell type-specific manner and in the context of physiological conditions at organismal level. The present study starts to address this gap by engineering the RanBP9-TurnX mouse.

The generation of this tool was possible because of the previous engineering of the RanBP9-TT model, which clearly showed that the addition of small tags at the C-terminus of RanBP9 enabled the unequivocal identification of the protein but did not cause any deleterious effect or alterations of RanBP9 interactions ^21^. Like RanBP9-TT mice, also the RanBP9-TurnX homozygous mice do not show lethality and sterility observed in RanBP9 null animals, nor any other obvious phenotype (***Figure 1***, ***Supplementary Figure 1***) ^7, 34^.

The design of the TurnX allele mimicked the Cre-mediated Recombination-Induced Tag Exchange (RITE) strategy used in lower organisms ^71^. More recently, the RITE approach was used also in human cells ^72, 73^. However, the RanBP9 TurnX model represents the first example of RITE in mouse. Notably, the TurnX allele that we designed does not include selection cassettes that were necessary to generate RITE alleles in human cells and lower organisms (***Figure 1***, ***Supplementary Figure 1***) ^71–73^. All considered, the TurnX mouse is relatively easy to engineer by CRISPR targeting and represents a prototype that can be mimicked to create other *in vivo* models when the addition of small tags does not interfere with the biological functions of the protein of interest.

The RanBP9-TurnX mouse can be used for countless studies where it can be one “module” in compound models studying pathological conditions where RanBP9 and the CTLH complex might have an important role such as neurodegenerative diseases and cancer ^1, 5, 10, 34, 74^. For example, RanBP9-TurnX mice can be coupled with compound transgenic models of Alzheimer’s disease ^75^ that include neuron-specific Cre ^76^, and investigate the interactions that RanBP9-V5 entertains in neurons during the course of the disease. Specifically, it could be useful to validate and follow the appearance of a truncated form that had been reported to arise in Alzheimer’s models ^77^. In the context of cancer, multiple permutations will be possible. One combination could be using a Cre-driven tumorigenesis model (like a lox-STOP-lox models, for example) ^78^ where cancer cells will produce RanBP9-V5 while the rest of the non-tumoral cells will display RanBP9-3xHA. Alternatively, more sophisticated, but extremely valuable, models can include Flpe-driven tumorigenesis in combination with cell type specific Cre lines to study RanBP9-interactions in the tumor microenvironment. The existence of inducible Cre lines might also allow to study the dynamic changes of interactions at different stages of tumorigenesis.

The RanBP9-TurnX model will be invaluable also for investigations to elucidate the role of RanBP9 and the CTLH complex in the regulation of physiological processes like metabolism and the immune response, two areas where current anecdoctal evidence suggests an involvement of this intriguing protein ^43, 79^.

In regard to metabolic investigations, the RanBP9-TurnX model can finally enable experiments to elucidate in the context of the whole organism whether RanBP9 and the CTLH complex interact with specific enzymes in basal or nutrient-restricted conditions. Importantly, Gid1/Vid30, the yeast ortholog of RanBP9, is key for the disposal of futile gluconeogenesis enzymes ^80^. The RanBP9-TurnX model offers the opportunity to perform studies *in vivo* aimed at establishing whether the CTLH complex binds and regulates gluconeogenesis in mammalian cells in organs like kidney and liver where this pathway is mainly enacted ^81^. In this regard, we validated the functionality of the TurnX allele in all the tissues we analyzed (***Figure 2***, ***Supplementary Figure 2***). Using a Cre ubiquitously expressed, we demonstrated that the licensing of RanBP9 tagged with V5 is accomplished in lung, liver, kidney, spleen, and thymus. RanBP9-V5 is detectable even in organs where the expression of RanBP9 is relatively low like the liver (***Figure 2C-G***). RanBP9-V5 is clearly detected both in kidney and liver where it can be efficiently immunoprecipitated with its CTLH binding partners (***Figure 2***, ***Figure 3***).

After confirming that the presence of Cre *in vivo* turned RanBP9-3xHA into RanBP9-V5, here we used the RanBP9-TurnX model to begin to investigate RanBP9 interactions in lung macrophages. This specific interest was prompted by the earlier observation that RanBP9 is highly expressed in lung alveolar macrophages, both in human and mouse, both in normal conditions and in tumor associated macrophages of non-small cell lung cancer masses ^9, 10, 12, 21^. To accomplish our goal, we mated the RanBP9-TurnX model to the LysM-Cre mouse, a strain commonly employed to study macrophage biology, which presents some limitations ^24^. Although it is the only strain available that expresses Cre in nearly all the macrophages in the airways, regrettably, LysM-Cre mice show Cre expression also in a variable percentage of neutrophils, classical dendritic cells, and some lung epithelial cells ^25^. Therefore, the LysMBP9X compound strain used for this study did not allow us to obtain the expression of a Cre-recombined RanBP9-V5 that is fully specific of macrophages, but only highly enriched. Notwithstanding these limitations, the LysMBP9X compound model enabled polished proteomic studies and reveled new interactions that the CTLH complex has in physiological conditions in lung.

We chose an approach where immunoprecipitation of RanBP9 from whole lung lysates is followed by mass spectrometry because it is an especially sensitive technology that can be powerful when appropriate negative controls are used. The presence of the HA and the V5 tags in the LysMBP9X mice allows the purging of the results from non-specific bindings due to similarities with the tags themselves using LysBP9 WT controls. This extra layer of negative controls cannot be included when employing commercially available antibodies recognizing RanBP9.

We established that the fulfillment of at least three key conditions needed to be met in order for the proteomic data of this lung macrophage-focused investigation to be meaningful. First, the immunoprecipitation both HA and V5 should pull down the core of known RanBP9 interactors. Second, the immunoprecipitation by RanBP9-3xHA and RanBP9-V5 should yield significantly different lists of proteins. Third, the immunoprecipitation using RanBP9-V5 should produce a proteome enriched in proteins expressed in myeloid/macrophagic cells.

The first condition was successfully met because the immunoprecipitation by both RanBP9-3xHA and RanBP9-V5 pulled-down from the same lung lysates all the known members of the CTLH complex. The only notable exception was Rmnd5b that was not significantly detected in the RanBP9-V5-associated proteome. This observation raises the possibility that this CTLH member, which is mutually exclusive with its paralog Rmnd5a in the formation of the multi-subunit complex ^13^, may be expressed at very low levels, if any, by LysM-Cre^POS^ cells including macrophages. The second and more challenging condition was also met because more than 90% of the proteins immunoprecipitated by RanBP9-3xHA were different in comparison to those immunoprecipitated by RanBP9-V5 from the same lung lysates (***Table 2***, ***Table 3***, ***Supplementary Table 1-4***). Ultimately, the fulfillment of the first two conditions clearly indicated that the model worked as designed and can discriminate RanBP9 interactions in a cell type specific manner.

The third condition that we had set was also fulfilled because the proteome immunoprecipitated by RanBP9-V5 includes proteins that are expressed in myeloid cells or involved in the innate immune response not pulled down by RanBP9-3xHA. For example, RanBP9-V5 co-immunoprecipitated Lyz2 and Msr1 (CD204) that are typically expressed by monocytes/macrophages. The presence of these two proteins alone strongly supports the successful enrichment of a myeloid cell specific proteome immunoprecipitated by RanBP9-V5. However, RanBP9-V5 immunoprecipitated with additional proteins with established roles in macrophages such as Anxa5, B2m, Cux1, Prdx5, and Tmed7 (***Table 3***, ***Supplementary Table 4***). These results clearly lay the groundwork for the mechanistic investigations of the role of RanBP9 in macrophage functions through these newly discovered cell-specific interactions.

In this regard, the rough analysis of all the interactors immunoprecipitated by both RanBP9-3xHA and RanBP9-V5 using the publicly available Metascape tool ^26^ clearly showed that a high number of RanBP9-interactions are with proteins involved in RNA processing at various levels (***Figure 5***). Therefore, RanBP9 and the CTLH complex may regulate RNA processing at different levels from transcription to translation. This is in agreement with previously published evidence ^19, 20, 27, 28^. A number of RNA transcription, RNA splicing, and RNA processing interactors identified here had not been reported before and illustrate how pervasive the modulation of RNA production, quality, and turnover by the CTLH complex could be. Since this E3 ligase appear to respond to various types of cell stress ^5^, it can be hypothesized that one consequence of CTLH action is a profound change of the RNAome upon stress.

Interestingly, both the RanBP9-3xHA and the RanBP9-v5 interactomes included proteins with an established role in the DDR that not only can help explain previous findings, but implicate RanBP9 in DNA repair processes other than homologous recombination (***Table 2***, ***Table 3***, ***Supplementary Table 2***, ***Supplementary Table 4***). For example, it is tempting to speculate that the interaction with Tiprl is instrumental to RanBP9 effects on the ATM-p53-γH2AX signaling axis ^9, 36, 37^. On the other hand, RanBP9-V5 from LysM-Cre^POS^ cells pulled down Prkdc and Atrx, that are both involved in triggering the innate immunity response ^67, 68^.

One area where the RanBP9 can be extremely valuable is in the investigation is the role of the regulation of cell and organismal metabolism and bioenergetics. Although RanBP9 has been suggested to regulate glycolysis in mammalian cells and carbon metabolism in yeast, results from this investigation and other anecdotal evidence suggests that the CTLH complex interact with enzymes that are involved in biosynthetic pathways other than just glycolysis. In this regard, a more detailed analysis of the RanBP9-TurnX proteome, brought to light interesting facts.

Several enzymes and proteins governing cell metabolism were present both in the RanBP9-3xHA and the RanBP9-V5 immunoprecipitated proteomes (***Table 2***, ***Table 3***, ***Supplementary Table 2***, ***Supplementary Table 4***). However, glycolytic enzymes were enriched only in the LysM-Cre^POS^ fraction. This result suggests that maybe the CTLH complex regulates glycolysis particularly in cancer cells where this enzymatic pathway is enhanced, or normal cells such as macrophages where glucose utilization is central to their polarization and function ^69, 70^.

Both the RanBP9-3xHA- and RanBP9-V5-associated proteomes included interactors that are involved in nucleotide, amino acid, and lipid metabolism as previously reported in other studies (***Table 2***, ***Table 3***, ***Supplementary Table 2***, ***Supplementary Table 4***) ^43, 48, 82^. Therefore, the CTLH complex might exert a pervasive regulation of cell metabolism. In this regard, it is also interesting that RanBP9 pulled down Rictor (previously reported RanBP9 interactor) ^48^ and Prr5l member of the TOR complex 2, suggesting a functional link with this key metabolic signaling node.

Finally, we obtained the striking result of 33 mitochondrial proteins immunoprecipitated by RanBP9-3xHA (17 proteins, representing 6.4% of total excluding CTLH members) and RanBP9-V5 (18 proteins, constituting 9.4% of total excluding CTLH members) combined (***Table 2***, ***Table 3***, ***Supplementary Table 2***, ***Supplementary Table 4***). This is not the first proteomic study reporting mitochondrial proteins immunoprecipitated by CTLH proteins ^13, 21, 43, 48, 83–85^. However, to date, there is no evidence to date that RanBP9 and the CTLH complex are localized at or inside this organelle although, in stressed conditions, RanBP9 was shown to migrate to the mitochondria and mediate apoptosis in concert with p73 ^86, 87^. In contrast with current knowledge, the present results strongly suggest that RanBP9 may be localized at the mitochondria in normal physiological conditions. It is also striking that of the 17 and 18 mitochondrial proteins immunoprecipitated by RanBP9-3xHA and RanBP9-V5 respectively, only two (2) were in common. This may suggest a role for RanBP9 and the CTLH complex that is “mitochondrial-specific”, where the mitochondria from LysM-Cre^POS^ cells are biochemically and functionally different from the mitochondria of the LysM-Cre^NEG^ cells. Future studies will be required to explore a possible role of RanBP9 in connection with mitochondrial functions.

In summary, we presented here an innovative tool to elucidate the cell type specific interactions that RanBP9 entertains *in vivo* both in physiological and pathological conditions. Results demonstrated that the RanBP9-TurnX model works as designed and provided the first report of a comprehensive LysM-Cre^POS^ -enriched RanBP9-interactome in normal lung cells that includes previously unreported RanBP9 interactions. This is not surprising since this is the first study performed *in vivo*, in normal lung cells, and in non-stressed conditions. The Cre-recombined RanBP9-TurnX model revealed a lung proteome with enrichment of interactions in LysM-Cre^POS^ cells listing proteins that are mostly expressed in macrophages. Also, this work establishes the RanBP9-TurnX model as a powerful tool to study the dynamic interactions of RanBP9 and the RanBP9-built CTLH complex in different cell types in the context of the whole organism.

## Acknowledgments

The authors express their gratitude to Dr. Tsonwin Hai for critically discussing findings, reading, and editing the manuscript. The authors acknowledge the contribution to this work of the Gene Editing Shared Resource, the Genetically Engineered Mouse Modeling Core, and the Genome Sequencing Shared Resource supported by grant the P30 CA016058 grant. This work was supported by the NIH R03CA259389 to V.C. and by the Pelotonia Idea Award GR123713 “Establishing the role of CTLH proteins in NSCLC” to V.C. This work was also supported by the Pelotonia Institute of Immuno-Oncology (PIIO). The content is solely the responsibility of the authors and does not necessarily represent the official views of the PIIO. Figure 1A, Figure 2A, and Supplementary Figure 1A were made using Biorender (www.biorender.com).

## Author Contributions

Designed the research study: D.P., and V.C.; conducted the experiments: Y.K., An.Te., A.O., Al.Th., M.C.C., F.D., F.A., M.C., S.H.A.S., L.R., L.Z., J.M.A., A.A., and V.C.; provided resources: D.P.C., G.F., V.C.; acquired the data: Y.K., An.Te., Al.Th., L.Z.; analyzed the data: Y.K., L.Z., M.F., P.D., and V.C.; wrote and/or edited the manuscript: Y.K., An.Te., J.M.A., D.P.C., A.L., G.F., M.F., D.P., and V.C.; project conception: D.P. and V.C.

## Data availability

The primary proteomic data of this manuscript are deposited and available upon request to the corresponding author at: ftp://MSV000093949@massive.ucsd.edu

---

**Supplementary Figure 1.**
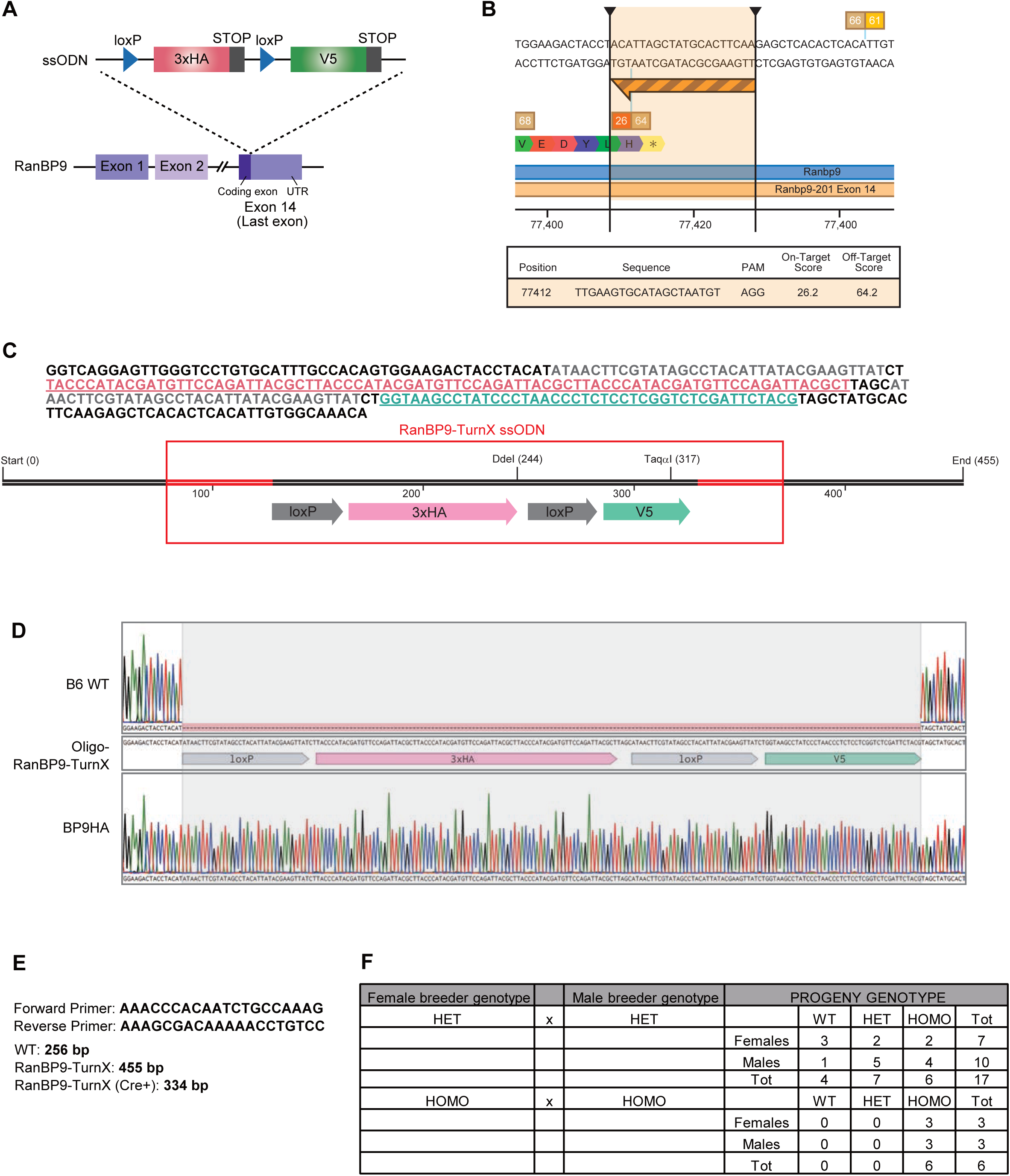
Generation of the RanBP9-TurnX allele. **A**) The 3x copies of HA are inserted right before the RanBP9 STOP codon of the last coding exon. **B**) Guide RNA used to target RanBP9 insertion site (same as Soliman *et al.*, 2020). **C**) Sequence of the ssODN used to engineer the RanBP9-TurnX allele. **D**) Sanger sequencing of RanBP9-Turnx homozygous animals confirms the correct insertion of the loxP-3xHA-loxP-V5 cassette as designed. **E**) Forward and reverse primers used to genotype RanBP9-TurnX animals. They amplify a product of 256 bp from the RanBP9 WT allele and a 455 bp product form the RanBP9-TurnX allele, respectively. **F**) RanBP9-TurnX heterozygous parents produce the expected numbers of WT, heterozygous, and homozygous male and female animals born from RanBP9 heterozygous parents. RanBP9-TurnX homozygous animals are fertile and do not show any obvious phenotype.

**Supplementary Figure 2.**
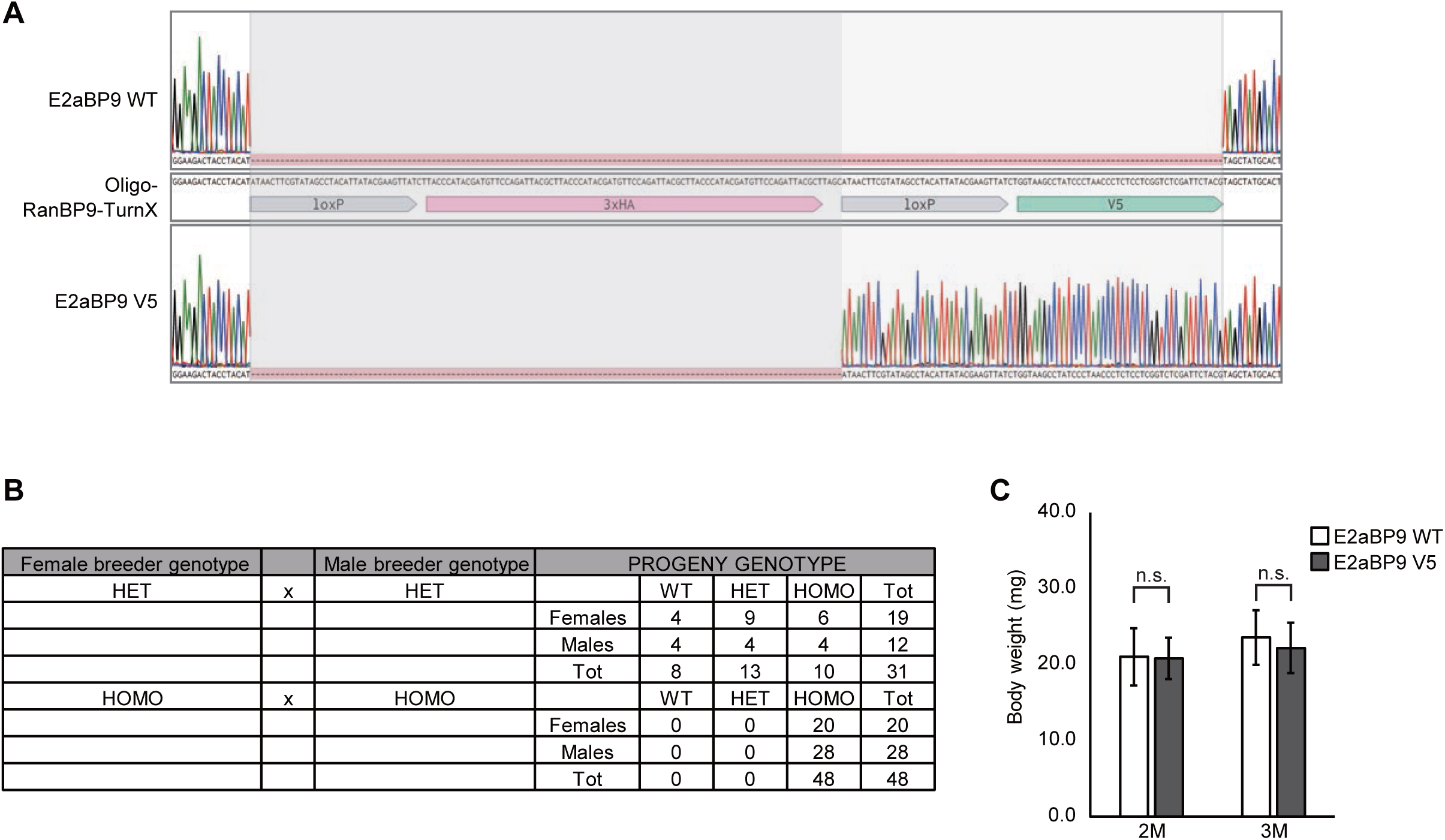
Cre-lox recombination in RanBP9-TurnX mice turns RanBP9-3xHA into RanBP9-V5. A) Sanger sequencing of RanBP9-V5 turned homozygous animals confirms the excision of the loxP-3xHA-loxP cassette as designed. B) RanBP9-V5 heterozygous turned parents produce the expected numbers of WT, heterozygous, and homozygous animals with no gender bias. RanBP9-TurnX homozygous animals are fertile and do not show any obvious phenotype. C) RanBP9-V5 homozygous animals do not show any significant difference in body weight compared to WT animals measured at 2 (2M) and 3 (3M) months of age.

